# Phage display assisted discovery of a pH-dependent anti-α-cobratoxin antibody from a natural variable domain library

**DOI:** 10.1101/2023.05.08.539834

**Authors:** Tulika Tulika, Rasmus W. Pedersen, Charlotte Rimbault, Shirin Ahmadi, Line Ledsgaard, Markus-Frederik Bohn, Anne Ljungars, Bjørn G. Voldborg, Fulgencio Ruso-Julve, Jan Terje Andersen, Andreas H. Laustsen

## Abstract

Recycling antibodies can bind to their target antigen at neutral pH in the blood stream and release them upon endocytosis when pH levels drop, allowing the antibodies to be recycled into circulation via FcRn-mediated pathway, while the antigens undergo lysosomal degradation. This enables recycling antibodies to achieve the same therapeutic effect at lower doses than their non-recyclable counterparts. The development of such antibodies is typically achieved by histidine doping of the variable regions of specific antibodies or by performing *in vitro* antibody selection campaigns utilizing histidine doped libraries. While often successful, these strategies may introduce sequence liabilities, as they often involve mutations that may render the resultant antibodies to be non-natural. Here, we present a methodology that employs a naïve antibody phage display library, consisting of natural variable domains, to discover antibodies that bind α-cobratoxin from the venom of *Naja kaouthia* in a pH-dependent manner. Upon screening of the discovered antibodies with immunoassays and bio-layer interferometry, a pH-dependent antibody was discovered that exhibits an 8-fold higher dissociation rate at pH 5.5 than 7.4. Interestingly, the variable domains of the pH-dependent antibody were found to be entirely devoid of histidines, demonstrating that pH-dependency may not always be driven by this amino acid. Further, given the high diversity available in a naïve antibody library, the methodology presented here can likely be applied to discover pH-dependent antibodies against different targets *ab initio* without the need of histidine doping.

**For broader audience:** Here, we present the discovery of an α-cobratoxin targeting pH-dependent antibody, with a variable region devoid of histidines, from a naïve antibody library with natural variable domains. Our findings suggest that the commonly taken approach of histidine doping to find pH-dependent antibodies may not always be required, and thus offer an alternative strategy for the discovery of pH-dependent antibodies.

## Introduction

Monoclonal antibodies (mAbs) of the immunoglobulin G (IgG) class are a rapidly growing class of drugs^1^ used to treat a range of conditions including cancer and autoimmune diseases^2,3^. The major factors behind their clinical success, include their high specificity and affinity for cognate antigens combined with the ability to mediate effector functions^2,3^. In addition, IgG has a plasma half-life of 3 weeks on average in humans, which makes it an attractive choice for the development of mAbs for diseases where exposure over time is key. This hallmark is regulated by binding of the IgG fragment crystallizable (Fc) region to a broadly expressed cellular receptor named the neonatal Fc receptor (FcRn), which rescues IgGs from intracellular lysosomal degradation via recycling or transcytosis. Mechanistically, this happens in a strictly pH-dependent manner where IgG is entering cells via fluid-phase pinocytosis followed by engagement of FcRn, which predominantly resides in mildly acidified endosomes. The complex is then recycled back to the cell surface or transcytosed across polarized cells followed by exposure to the near neutral pH, which triggers dissociation and release of the IgG to the extracellular milieu^4,5^. As such, IgG antibodies are rescued from intracellular degradation via FcRn-directed transport routes.

However, when most IgGs are bound to their cognate antigen, this often occurs with high affinity throughout this endosomal pH gradient. Thus, the IgGs may undergo antigen-mediated clearance via lysosomal degradation or they may be recycled by FcRn along with the bound antigen. As a result, antibodies can bind the antigen only once in their lifetime. An attractive strategy is to engineer antibody binding to the antigen such that high affinity is kept at near neutral pH while binding becomes weaker when approaching the acidic environment of endosomes^6–15^. This allows the antigen to dissociate from the antibody in the acidic endosomes (∼pH 5.0-6.5) and undergo lysosomes for degradation, while the antibody is rescued via FcRn-mediated pathway and released upon exposure to the near-neutral pH conditions (pH ∼7.4) at the cell surface. This will allow the same IgG to be used multiple times, ready to engage new antigens in the blood stream^6,13,16^. Such engineering has shown to reduce the required dose and/or frequency of dosing to achieve therapeutic effect^14,17^. Importantly, this is an attractive approach for design of antibodies tailored for treatment regiments relying on high dosing and where cost is a limiting factor, such as snakebite envenoming and infectious diseases^18^.

Specifically, the ability of antibodies to bind cognate antigens in a pH-dependent manner has largely been attributed to the presence of histidines (p*Ka* ∼6.0) at the antibody-antigen binding interface^19,20^. Thus, the discovery of pH-dependent antibodies has predominantly been carried out using histidine scanning approaches or histidine-enriched libraries^8–12,21–26^. However, such strategies may not always be straightforward as histidine doping may compromise target binding properties at neutral pH^12,23,24^. In addition, histidine-mediated pH-dependent binding requires the epitope to have positively charged residues, which restricts the number of suitable epitopes^27^. Furthermore, histidine-enriched antibody libraries are designed, and therefore, the generated antibodies may have developability and immunogenicity risks that should be taken into consideration. Immunized libraries have also been explored but derived antibodies have needed to undergo humanization^28,29^. Additional approaches for discovery of pH-dependent antibodies are thus attractive.

In this study, we demonstrate the utility of natural naïve human antibody libraries for discovery of fully human IgG1 antibodies with pH-dependent antigen binding properties. We show that this is possible even without having any histidines in the complementarity determining regions (CDRs). To achieve this, we employed phage display technology based on a naïve human antibody library consisting of naturally occurring variable domains for the discovery of pH-dependent antibodies against a long-chain α-neurotoxin, namely α-cobratoxin (α-cbtx). The results showcase that pH-dependency is not always driven by histidines, but instead can be due to other structural features at the antibody-antigen binding interface.

## Results

### pH elution during antibody phage display selections enables discovery of pH-dependent binders

To enrich antibodies with a pH-dependent antigen binding, three consecutive rounds of phage display selections were performed against biotinylated α-cbtx using a buffer with low pH or trypsin (as a control) for elution of binding single-chain variable fragment (scFv) displayed on phages. Following reformatting to soluble scFv and expression in *E*.*coli*, 918 of the 1472 screened clones bound to α-cbtx in an expression-normalized capture (ENC) dissociation-enhanced lanthanide fluorescence immunoassay (DELFIA)^30,31^ with a signal above the arbitrary cut-off value of 5000 (Figure 1(A)). To screen for pH dependent binding, 635 of the binding clones were randomly selected, re-expressed, and analyzed in an ENC pH DELFIA, where the clones were allowed to bind α-cbtx at pH 7.4 and thereafter either incubated in a buffer of pH 7.4 or pH 5.4 for an hour before adding the detection reagent. This revealed that 166 clones showed at least 50% decrease in the binding signal after incubation at pH 5.5 compared to 7.4, indicating a pH-dependent binding of the scFvs to the antigen (Figure 1(B))^8^. Further, sequencing of these 166 clones showed that ∼99% of the clones were identical, resulting in 2 unique clones. Both clones came from the phage display selection where a low pH buffer was employed for elution of the bound phages.

**Figure 1:**
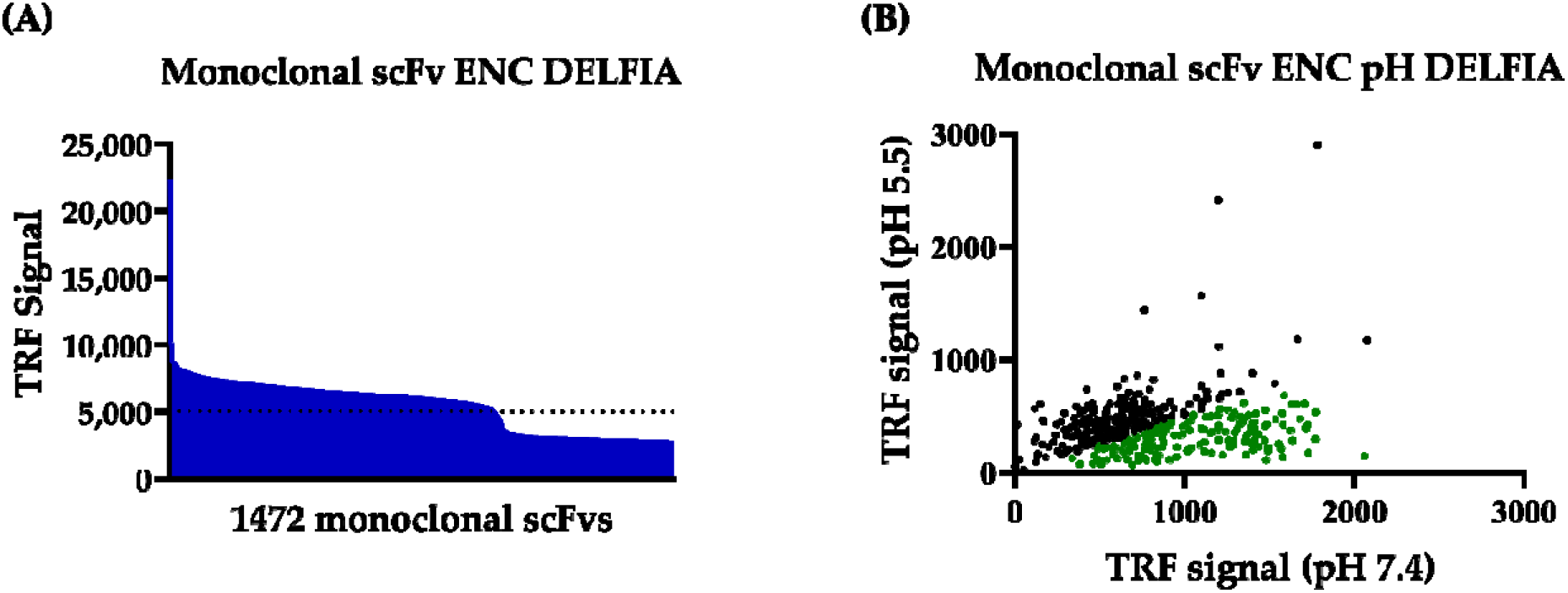
Screening of binders. **(A)** Binding signal of 1472 discovered monoclonal single-chain variable fragments (scFvs) against α-cobratoxin (α-cbtx) in expression-normalized capture (ENC) dissociation-enhanced lanthanoid-based fluorescence assay (DELFIA). **(B)** Scatter plot showing binding signal of a subset of selected α-cbtx binding monoclonal scFvs ENC pH DELFIA. The monoclonal scFvs showing at least 50% decrease in binding signal after being incubated at pH 5.5 than pH 7.4 buffer are coloured in green.

### The most abundant pH-dependent clone contains no histidine residues in the variable domains

The two unique scFv clones were expressed in *E. coli*, and the bacterial supernatants containing the expressed scFvs were used in bio-layer interferometry (BLI) to determine their pH dependent target dissociation. The most abundant clone, TPL0197_01_C08 (which will be referred to as C08 from here on) showed a faster off-rate at pH 5.5 compared to pH 7.4 (Figure 2(A)). Sequence analysis of this clone showed that it contains no histidines in the variable regions-neither in the CDRs nor in the framework (Figure 2(B)). This was surprising since histidines have been widely attributed as a major contributing factor of pH-dependence^6^.

**Figure 2:**
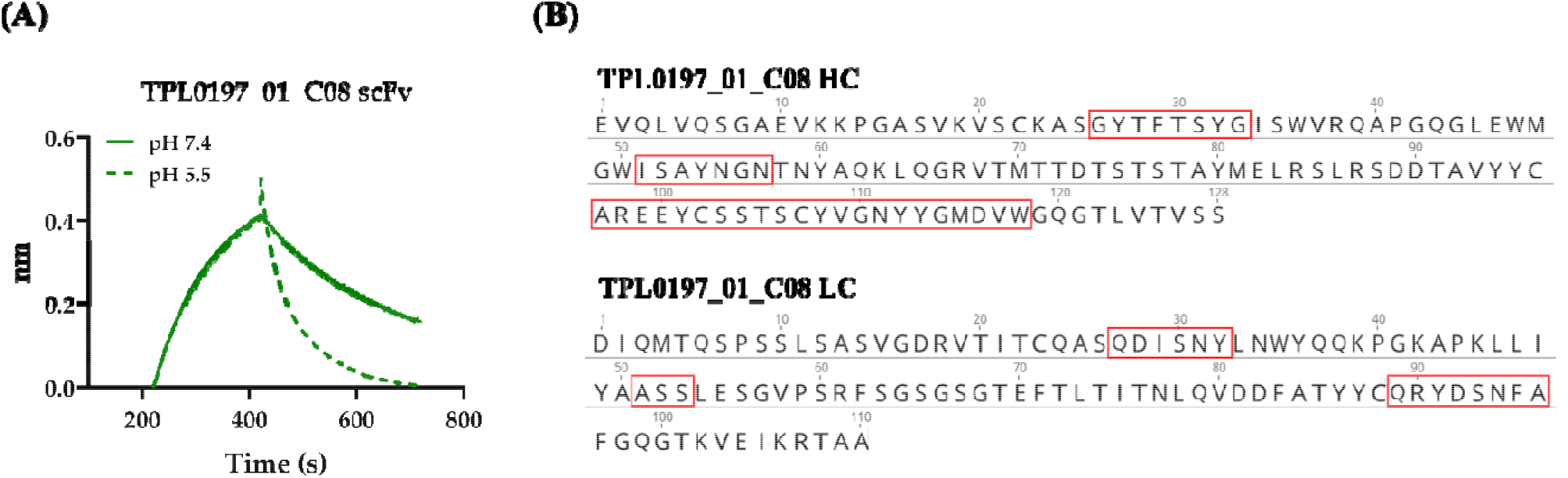
Bio-layer interferometry (BLI) binding curves and sequence of the pH-dependent clone C08. **(A)** BLI sensogram showing association at pH 7.4 and dissociation at pH 7.4 (solid line) and 5.5 (dashed line for scFv C08). **(B)** Amino acid sequences of heavy chain (HC) and light chain (LC) of clone C08. The CDRs are highlighted in red boxes.

### The pH dependent IgG, C08, shows less binding at pH 5.5 compared to pH 7.4 in ELISA

To further characterize C08, it was reformatted to an IgG1 format, expressed in CHO cells, and purified. To validate IgG pH-dependent binding to α-cbtx, an ELISA-based binding assay was performed at pH 7.4 and 5.5 (Figure 3(A)). A high affinity (1.8 nM), α-cbtx targeting IgG 2554_01_D11 (which will be referred to as D11 in the following text) that previously discovered through phage display without any pH-selection pressure, was included was for comparison^32^. Both IgGs bound to α-cbtx at pH 7.4, however the signal at comparative concentrations was lower for C08 than D11, indicating a low affinity of C08 towards α-cbtx. At pH 5.5, C08 showed negligible binding to α-cbtx, while D11 bound with almost identical strength to α-cbtx as that observed at pH 7.4 (Figure 3(A), (B)). This showed that both binding and dependency of C08 towards a-cbtx was retained after reformatting to an IgG format.

**Figure 3:**
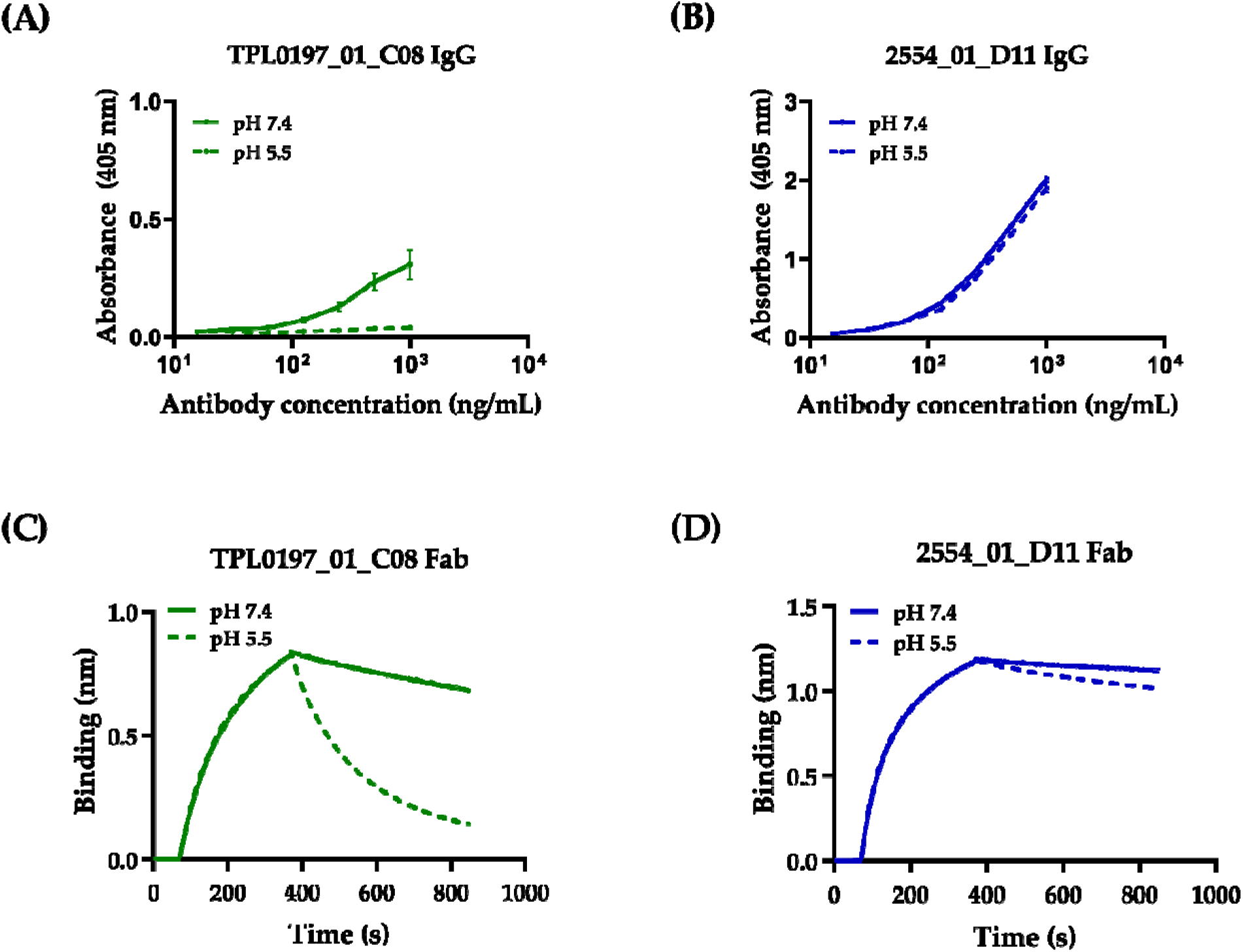
Binding characterization of a pH-dependent and a non-pH-dependent antigen binding clone in IgG and Fab format. ELISA binding curves to α-cobratoxin (α-cbtx) at pH 7.4 (solid line) and 5.5 (dashed line) of **(A)** pH-dependent IgG C08 and **(B)** non-pH dependent IgG D11. Bio-layer interferometry (BLI) curves showing association at pH 7.4 and dissociation at pH 7.4 (solid line) and 5.5 (dashed line) for a **(C)** pH-dependent Fab C08 and **(D)** non-pH dependent Fab D11.

### pH-dependent antigen binding Fab C08 shows increased rate of dissociation at pH 5.5

To determine the binding kinetics of the pH-dependent binder C08 while avoiding avidity effects, C08 was reformatted into a Fab, expressed in CHO cells, and His-tag purified. D11 in Fab format was included as a non-pH-dependent antigen binding control. Both Fabs were then characterized for their pH-dependent binding using BLI, on biotinylated α-cbtx loaded streptavidin biosensor tips. Fabs were allowed to associate at pH 7.4 followed by dissociation at either pH 7.4 or 5.5 (Figure 3(C), (D)). An 8-fold higher dissociation rate at pH 5.5 than 7.4 was observed for C08. In contrast, the difference between the dissociation rates at these pH values was 1.5-fold for D11 (Table 1). Thus, the pH-dependent binding of C08 towards α-cbtx was confirmed in both IgG and Fab formats using ELISA and BLI respectively.

**Table 1:** Dissociation rates of anti-α-cbtx Fabs

| Fab | $k_d$ (s<sup>-1</sup>)<br>pH 7.4 | $k_d$ (s<sup>-1</sup>)<br>pH 5.5 | Fold difference<br>(pH 5.5/7.4) |
| --- | --- | --- | --- |
| TPL0197_01_C08 | $8.077 \times 10^{-4}$ | $6.634 \times 10^{-3}$ | 8.22 |
| 2554_01_D11 | $2.61 \times 10^{-4}$ | $4.01 \times 10^{-4}$ | 1.53 |

Dissociation rates of pH-dependent and non-pH-dependent Fab binders determined by BLI. The association of Fabs to the biotinylated α-cbtx loaded on streptavidin biosensors was performed at pH 7.4, while the dissociation was performed at pH 5.5.

## Discussion

In recent years, recycling antibodies i.e., antibodies that engage with their target antigen in a pH-dependent manner have gained increased attention, as they retain high efficacy when administered at lower doses than their non-pH-dependent counterparts. This feature stems from the combination of pH-dependent antigen binding properties and long plasma half-life deriving from FcRn-mediated recycling mechanism. Consequently, various strategies to discover and engineer pH-dependent antigen-binding antibodies have been explored^7^.

In this study, we employed a naïve human antibody phage display library consisting of naturally occurring variable domains and successfully discovered an antibody that binds its target antigen in a pH-dependent manner. While pH-dependent antibodies have previously been discovered using histidine doping of antibody libraries or the paratopes of a pre-existing antibodies^7^, we here show that such strategies may not necessarily be required to discover antibodies with highly pH-dependent antigen binding properties. Avoiding the incorporation of histidines in variable regions of antibodies may have several benefits. First, introduction of such mutations in antibody paratopes can be a time-consuming and laborious task, as it requires detailed analysis and validation of pH-dependency of the mutants and may still not provide an antibody with desirable features. For example, it has been reported that, occasionally, histidine mutations that were incorporated to reduce the binding between antibody and antigen at low pH also resulted in reduced binding at neutral pH^7,12^. Second, the incorporation of mutations, particularly in antibody sequences deriving from natural sources, may come with a risk of introducing sequence liabilities in the antibody that could pose problems for the developability of the antibody, such as causing it to be immunogenic^33–36^. The employment of an antibody library with natural variable domains for the discovery of pH-dependent antibodies could potentially reduce this risk, since the variable domains of the antibodies have gone through the test for self-tolerance^37^. Third, given that a naïve natural antibody library can be used to find binders to a multitude of targets, the observation that histidine doping may not be necessary for the discovery of antibodies with pH-dependent antigen-binding properties indicates that such antibodies can possibly be found against many targets directly *ab initio*^38^.

Besides originating from a naïve natural antibody library, the variable domains of the discovered pH-dependent antibody were surprisingly found to be entirely devoid of histidine residues. Although the presence of histidine is widely attributed to mediate pH-dependent binding^28^, our findings suggest that antibodies can, at least at times, derive pH-dependent antigen binding properties from different residues. A possible explanation for this could be the effect of the molecular microenvironment that can change the local pKa of amino acids at the antibody-antigen binding interface, which may influence the binding behavior between the two molecules^39,40^.

So far, the utility of recycling antibodies that bind their target antigens in a pH-dependent manner has mostly been demonstrated against endogenous targets, such as IL-6, PCK9, CXCL10, and TNF-α ^8–11^. In this study, an exogenous soluble antigen, α-cbtx from *N. kaouthia* venom, was used as the target antigen. Recycling antibodies that bind snake toxins in a pH-dependent manner could potentially find utility for the development of novel types of antivenoms, which could be administered to patients at lower doses compared to both current plasma-derived antivenoms and recombinant antivenoms which are based on non-recyclable antibodies. However, in the case of snakebite envenoming, both complex toxicokinetics and pharmacokinetics are at play^41^. While endogenous targets are often (semi-) constitutively produced within the body and therefore can be maintained at a concentration below certain thresholds by using recycling antibodies; toxins are instantaneously injected in a large dose into the body of the victim during a snakebite envenoming case, and thus, require an urgent intervention^18,41^. The effectiveness of recycling antibodies with pH-dependent binding to their target under such circumstances, where large amounts and often fast-acting toxins are required to be removed from circulation is not known. However, given that, in the majority of the cases of snakebite envenoming, the bite is either intramuscular or subcutaneous, the injection of toxins is followed by an initial absorption phase^41^. Further, for systemic toxins (such as α-cbtx), a likely depot effect might result in a delay in the onset of their toxic effects since the toxins first need to leave the bite site to enter the blood^18^. Under such circumstances, where the toxins are released over time from the bite site into circulation and not all at once, the antibodies would not be burdened with large amounts of toxins at once. Thus, we speculate that it would potentially be possible to neutralize the toxins using recycling antibodies at a lower dose and consequently at reduced cost, than the non-recyclable antibodies. However, to understand the overall effect of using recycling antibodies to neutralize toxins in a snakebite envenoming case requires further investigation. Nevertheless, the methodologies presented in this study could find broad applicability beyond snakebite envenoming as a general approach for the discovery of pH-dependent antibodies against potentially any target using *in vitro* display technologies.

## Materials and Methods

### Biotinylation of antigen

Purified α-cobratoxin (α-cbtx) from *N. kaouthia* (Latoxan, France) was dissolved in 1X standard phosphate buffered saline (PBS) and biotinylated using 1:1.25 (toxin: biotin reagent) molar ratio as previously described^30^. The biotinylated toxins were purified using buffer exchange columns (Vivacon 500, Sartorius, 3000 Da Molecular Weight Cut-Off) using the manufacturer’s protocol. Protein concentration was determined using the toxin’s extinction coefficient and absorbance measurement with a NanoDrop One instrument. The degree of biotinylation was analyzed by MALDI-TOF in an Ultraflex II TOF/TOF spectrometer (Bruker Daltonics).

### Solution-based phage display pH selection

The protocol for carrying out solution-based phage display selections was adapted from previous work^8^. The libraries used were the IONTAS naïve single-chain variable fragment (scFv) phage display κ library^37^. Briefly, the phage display library was first blocked using 3% skimmed milk in PBS (MPBS) and then deselected using streptavidin-coated beads (DynaBeads M280, Thermo Fisher #11205D). 100 nM biotinylated α-cbtx was mixed with the deselected library and incubated for selection for one hour. This selection was carried out at pH 7.4. Phages bound to α-cbtx were then captured on streptavidin-coated beads, and non-specific phages were eliminated by washing thrice with PBS + 0.1% Tween (PBS-T), and twice with PBS. In the first round of phage display panning, all phages were eluted by trypsin digestion. In round two and round three, pH-dependent clones were eluted by adding citrate buffer at pH 5.5 for 15-60 min or by trypsin. For the phages eluted using citrate buffer, trypsin was subsequently added to the eluted phages. The eluted phages were then used to infect TG1 cells as described before^42^.

### Sub-cloning and screening of scFvs

Sub-cloning of scFv genes from phage outputs into the pSANG10-3F vector and primary screening were performed as previously described^30^. In short, *NcoI* and *NotI* restriction endonucleases sites were used to sub-clone scFv genes from phagemids into the pSANG10-3F vector, which was then transformed into *E. coli* strain BL21(DE3) (New England Biolabs). From each of the selection outputs, 184 colonies were picked, expressed in 96 well format and assessed for binding to 50 nM of biotinylated α-cobratoxin in an expression-normalized capture (ENC) dissociation-enhanced lanthanide fluorescence immunoassay (DELFIA) as described earlier, with a few modifications^30^.

First, Nunc MaxiSorp plates (Invitrogen, 44-2404-21) were coated overnight with 50 μL of 2.5 μg/mL anti-FLAG M2 antibody (Sigma Aldrich, F1804). Plates were washed thrice with PBS and blocked with 200 μL of 3% MPBS. Plates were washed thrice with PBS, and 60 μL of 0.5X unpurified scFv-containing culture supernatant in 3% MPBS was added before incubating for 1 hour at room temperature. Plates were washed thrice with PBS + 0.1% Tween and thrice with PBS before adding 50 μL of 50 nM biotinylated α-cbtx in MPBS to each well. After 1 hour of incubation, the plates were washed thrice with PBS + 0.1% Tween and thrice with PBS. Then, 1 μg/mL of Europium-labeled Streptavidin (Perkin Elmer, 1244–360) in dissociation-enhanced lanthanide fluorescence immunoassay (DELFIA) Assay Buffer (Perkin Elmer, 4002–0010) was added. Following 30 minutes of incubation, plates were washed thrice with PBS + 0.1% Tween and thrice with PBS, and DELFIA Enhancement Solution (Perkin Elmer, 4001–0010) was added for detection of binding. Clones that gave a signal above 5,000 counts were selected for further characterization.

### ENC pH DELFIA and sequencing

To characterize the pH-dependency of the α-cbtx-binding scFv candidates, a modified ENC DELFIA assay was performed. The assay was carried out as described in section 2.3, with an additional pH-elution step. The ENC pH DELFIA was performed in duplicates until after the incubation and washing of α-cbtx, where 60 μL of citrate buffer at either pH 6.0 or pH 7.4 was added to each well. Following 60 minutes of incubation, plates were washed thrice with PBS + 0.1% Tween and thrice with PBS. Detection of biotinylated antigen was carried out as described in section 2.3.

### BLI off-rate screening of scFvs from bacterial culture supernatant

Prior to the assay, streptavidin (SAX) biosensors were pre-wetted for at least 10 min in 1x Kinetics Buffer (KB, Forte Bio). Screening assay was performed by first loading 1μg/mL biotinylated α-cbtx on SAX biosensors, followed by a 120 s baseline step in 1x PBS pH 7.4. The toxin-loaded biosensors were then dipped in scFv-containing bacterial supernatant wells for 600 seconds of association step, followed by a dissociation step in 1x PBS pH 7.4 for 600 seconds. The tips were regenerated in regeneration buffer 10mM Glycine pH 2.0 and neutralization buffer (1x KB) for 5 seconds for a total of 5 cycles. The tips were then dipped into 1x PBS pH 5.4 for 120 s, followed by association in the scFv-containing bacterial supernatant wells for 600 seconds, and dissociation in 1x PBS pH 5.4 for 600 seconds. The experiment was performed at 25°C with shaking at 1000 rpm. ForteBio’s data analysis software was used to fit the curves using a 1:1 binding model to obtain dissociation rates for pH 7.4 and 5.4.

### Kinetics measurements with Fab

The KD of the Fab was determined as described in 2.5, except that purified Fabs in 1x HEPES pH 7.4 were used for association at 200nM-3nM in a 2-fold dilution. ForteBio’s data analysis software was used to global fit the curves using 1:1 binding model to determine kinetic constants.

### Production of IgGs

The reformatting of scFv into IgG was performed as previously described^42^, except that the IgG expression vector for TPL0197_01_C08 contained the human kappa light chain. ExpiCHO cells were cultured and transfected with expression vector according to the manufacturer’s guidelines (Gibco™) following a protocol where ExpiFectamine™ CHO Enhancer and a single feed were added at Day 1 and cells were maintained at 37°C and 5% CO_2_. The supernatant was collected at Day 7 by removal of the cells through centrifugation at 300 g for 5 min, followed by an additional centrifugation at 1000 g for 5 minutes. The supernatant was either used for purification on the same day or stored at -80°C. Supernatant was thawed overnight at 4°C, centrifuged, filtered and loaded on a MabSelect column (Cytiva). 20 mM sodium phosphate and 150 mM NaCl (pH 7.2) was used for equilibration and washing of the column and elution was performed with 0.1 M sodium citrate (pH 3). Elution fractions were immediately neutralized by 1 M Tris (pH 9) using 1/5 V of neutralization solution for 1 V of elution fraction. Fractions of interest were pooled and loaded on a HiPrep 26/10 desalting column for buffer exchange to Dulbecco’s PBS. Protein fractions were sterile-filtered and concentrated by centrifugal filtration using an Amicon® Ultra-15 centrifugal filter unit (30 kDa NMWL). The final concentration was determined by measuring the absorbance at 280 nm on a Nanodrop2000 instrument. Purity was checked by SDS-PAGE. The purified protein was stored at 4°C or -80°C.

### Production of Fab

The reformatting of scFvs to Fabs and expression of Fabs was carried out as described in section 2.7 except that expression vector of TPL0197_01_C08 contained the constant domain 1 sequence of heavy and human kappa light chain, while that of 2554_01_D11 contained human lambda light chain.

After expression, the collected supernatant, centrifuged, and loaded on a 5-mL HisTrap Excel column (Cytiva), equilibrated with 20 mM sodium phosphate (pH 7.4), 500 mM NaCl. The column was washed with 10 column volumes of 10 mM imidazole in 20 mM sodium phosphate (pH 7.4), 500 mM NaCl. Elution was performed in up-flow mode with 20 column volumes of 500 mM imidazole in 20 mM sodium phosphate (pH 7.4), 500 mM NaCl. Protein containing fractions of interest were pooled and loaded on a HiPrep 26/10 desalting column for buffer exchange to Dulbecco’s PBS. The protein was then concentrated by centrifugal filtration using an Amicon® Ultra-15 centrifugal filter unit (10 kDa NMWL). The final concentration was determined by measuring the absorbance at 280 nm on a Nanodrop2000 instrument. Purity was checked by SDS-PAGE. The purified protein was stored at 4°C or -80°C.

### pH-dependent binding properties in ELISA

96-well EIA/RIA 3590 microplates (Corning) were coated with 100μL of 0.5μg/mL α-cbtx (Latoxan, France) diluted in PBS overnight at 4^°^ C. The plates were blocked with 4% skimmed milk powder (M) (Sigma-Aldrich) dissolved in PBS for 1 h, followed by washing four times with PBS containing 0.05% Tween20 (PBS-T) (Sigma-Aldrich) (PBST). Unless stated, the following steps were carried out at pH 7.4 and 5.5, respectively, and the washing was conducted with PBS-T with the corresponding adjusted pH. Next, 100μL of titrated amounts (1 – 0.015 μg/mL) of the samples containing IgGs diluted in PBST-M were added to the plates and incubated at RT for one hour. After washing, 100μL of ALP-conjugated anti-human IgG Fc-ALP (Sigma-Aldrich) diluted 1:5000 in M-PBST was added and incubating for 1 hour. Thereafter, following washing with PBST, the bound proteins were detected by adding 100μL of 1 mg/mL p-nitropenylphosphate substrate tablets dissolved in diethanolamine buffer (pH 9.8) (Sigma-Aldrich). The absorbance was measured at 405 nm using the Sunrise spectrophotometer (Tecan).

## Acknowledgments

The authors are supported by a grant from the European Research Council (ERC) under the European Union’s Horizon 2020 research and innovation programme (grant no. 850974); Villum Foundation under grant 00025302; Wellcome (221702/Z/20/Z); Novo Nordisk Foundation (NNF20SA0066621); and Research Council of Norway (287927). The authors would like to thank Sara Petersen Bjørn, Karen Kathrine Brøndum, and Daniel Duun from National Biologics Facility for the reformatting and production of Fabs and IgGs.

